# Visual and non-visual properties of filters manipulating short-wavelength light

**DOI:** 10.1101/596957

**Authors:** Manuel Spitschan, Rafael Lazar, Christian Cajochen

## Abstract

Optical filters and tints manipulating short-wavelength light (so-called “blue-blocking” or “blue-attenuating”) are used as a remedy for a range of ocular, retinal, neurological and psychiatric disorders. In many cases, the only available quantification of the optical effects of a given optical filter is the spectral transmittance, which specifies the amount of light transmitted as a function of wavelength. Here, we propose a novel physiologically relevant and retinally referenced framework for quantifying the visual and non-visual effects of these filters, incorporating the attenuation of luminance (luminance factor), the attenuation of melanopsin activation (melanopsin factor), the shift in colour, and the reduction of the colour gamut (gamut factor). We examined a novel data base of optical transmittance filters (n=120) which were digitally extracted from a variety of sources and find a large diversity in the alteration of visual and non-visual properties. We suggest that future studies and examinations of the physiological effects of optical filters quantify the visual and non-visual effects of the filters beyond the spectral transmittance, which will eventually aid in developing a mechanistic understanding of how different filters affect physiology.

## Introduction

Optical filters can be used to modify the visual input by blocking or attenuating light at specific parts of the visible spectrum^1,2^. So-called blue-blocking or blue-attenuating filters reduce the amount of short-wavelength light at the eye’s surface, the cornea. Optically, this is typically realised using one of two ways: 1) using a cut-off filter, which blocks or attenuates light below a specific wavelength, 2) using a notch filter, which blocks or attenuates light within a specific short-wavelength light.

Optical filters alter the spectral distribution of the light incident on the retinal surface relative to no filtering^3^. This has direct effects on the activation of the cones and rods in the retina, which allow us to see during the day and night, respectively. In addition, the activity of the melanopsin-containing retinal ganglion cells, is also modulated by the use of optical filters. These cells, though only making up <1% of retinal ganglion cells, are of great importance to non-visual functions, such as entrainment by the circadian rhythm to the environmental light-dark cycle, and the suppression of melatonin in response to light^4^.

Previous investigations have examined the effects of blue-blocking or -attenuating filters on visual performance^5-11^, colour vision^11-13^, and steady-state and dynamic parameters of pupil size^14,15^. Furthermore, they have received attention in the domain of sleep medicine and chronobiology, where their effects on melatonin suppression^16-23^, circadian rhythmicity^24-28^, sleep^6,15,19,23,27-31^, modulation of alertness by light^32,33^, and for use by shift or night workers^34,35^ have been investigated.

In addition, blue-blocking or -attenuating filters have been employed in the management of psychiatric and neurological (bipolar disorder^36-38^, depression^30^, ADHD^39^, blepharospasm^40,41^, migraine^42-44^, photosensitive epilepsy^45^, insomnia^30,39,46^) and retinal conditions (rod monochromacy/achromatopsia^47-50^, retinitis pigmentosa^51^, others^52^), as well as for reducing non-specific photophobia^44,53^ and eye fatigue^54^. Coloured filters have also been used previously to improve reading difficulties and relieve visual stress^55,56^ and symptoms of migraine^57^, though these are not specifically attenuating short-wavelength light^58^ (but may, depending on the chosen chromaticity).

Across these studies, filters by different manufacturers were used. The terms “blue-blocking” or “blue-attenuating” are not well-defined standardised and therefore, there is great ambiguity about the exact properties of filters carrying those names. The unique characteristic of a filter is its spectral transmittance, which is a quantified specification of how much light is passed through the filter at a given wavelength. But it is impossible to determine the precise effects on various visual and non-visual functions just by examining the spectral transmittance function of a filter. While the transmittance is necessary to determine these effects quantitatively, this cannot be done “by eye”.

By changing the spectral distribution of the transmitted light can have a range of different effects. Here, we considered four main effects (Fig. 1), all of which are important to assess the properties of a given filter:

1. *Luminance factor* [%]: By changing the spectral properties of the incident light, the luminance of the effective stimulus is altered. This attenuation can be expressed as the ratio between the luminance of the filtered spectral distribution to the luminance of the unfiltered spectral distribution. We call this the luminance factor. A luminance factor of 100% corresponds to no change in luminance by the filter. Values >100% are not possible.
2. *Melanopsin factor* [%]: Similarly, the activation of melanopsin may be reduced by the optical filter under investigation. Similar to the luminance factor, the melanopsin factor is expressed as the proportion of melanopsin activation with filter relative to without filter.
3. *Colour shift:* Optical filters with non-uniform transmittance also lead to a shift in the colour appearance of a colour which would appear as white without filter. With blue-blocking filters, these shifts typically occur in the yellow, orange or amber directions. We quantify this colour shift using a Euclidian distance metric in the uniform CIE 1976 u’_10_v’_10_ colour space based on the CIE 1964 10° observer.
4. *Gamut factor* [%]: Objects of different colours will appear different when seen with a filter. Without the filter, the distribution of differently coloured objects is called the gamut, which is simply the area which the object colours “inhabit” in a colour space. We can compare the area of the gamut when objects are seen through a filter with the area of the gamut without the filter.

**Figure 1:**
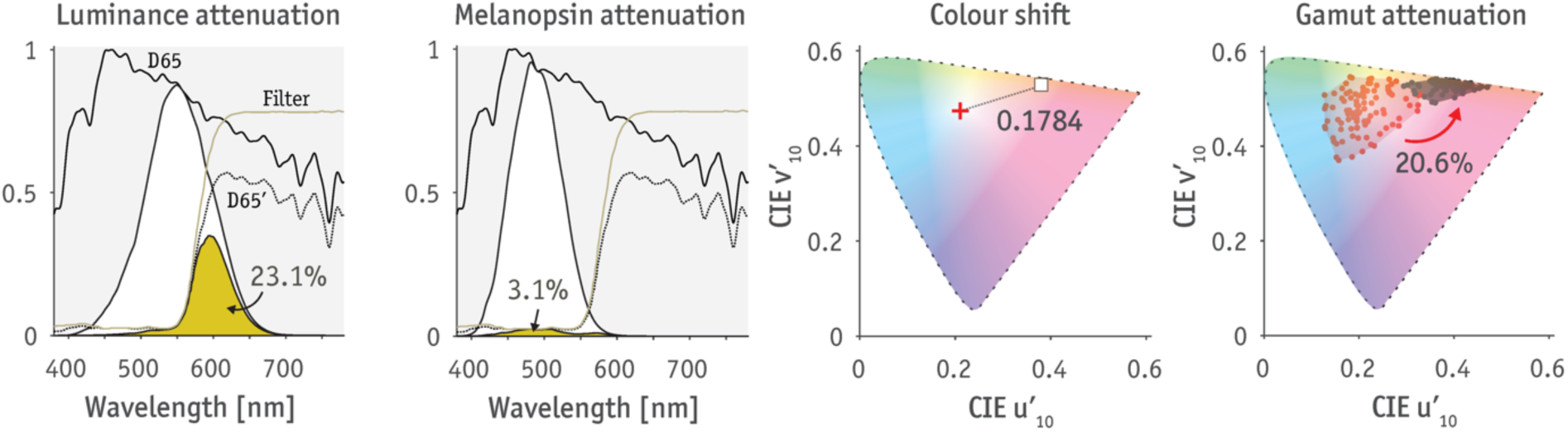
Visual effects of filters. Luminance factor is given as the ratio between the luminance of daylight seen with filter (shown as the beige area) to without filter (shown as the white area). In this case, the luminance with the filter is only about a quarter (23.1%), i.e. the light is only a quarter as bright. Melanopsin factor follows the same calculation, except for the melanopsin photopigment. Because the cut-off wavelength of the filter is outside of the spectral sensitivity of melanopsin, the attenuation is far larger compared to luminance (3.1%). Colour shift is the Euclidian distance in the uniform colour space used here. The smaller the number, the smaller the colour shift. Gamut factor refers to the reduction of the colour space which common surface reflectances inhabit, with and without the filter. In this case, the colour gamut with the filter is only around one fifth (20.1%) of the gamut without the filter. For demonstration purpose, the spectral power distribution was assumed that to be of noon daylight (D65, daylight with a correlated colour temperature [CCT] of 6500K).

We note that these four properties of filter are by no means exhaustive, but they allow for a strongly quantifiable and yet intuitive approach of the effects of different optical filters on visual and non-visual function. We also note that our analysis does not include consideration of transmission of light in the UV band and makes no claims about damaging effects of UV or other radiation. Similarly, we did not consider the polarization properties of the filters.

We started examining the visual and non-visual filters using simulated filters. We used an analytic description of the typical transmittance profile using a sigmoid function (Fig. 2). The model parameters were the (1) cut-off wavelength, (2) the upper asymptote, and (3) the slope of the function. By changing each of these parameters in isolation, we can examine how properties of the spectral transmittance functions affect the derived parameters.

**Figure 2:**
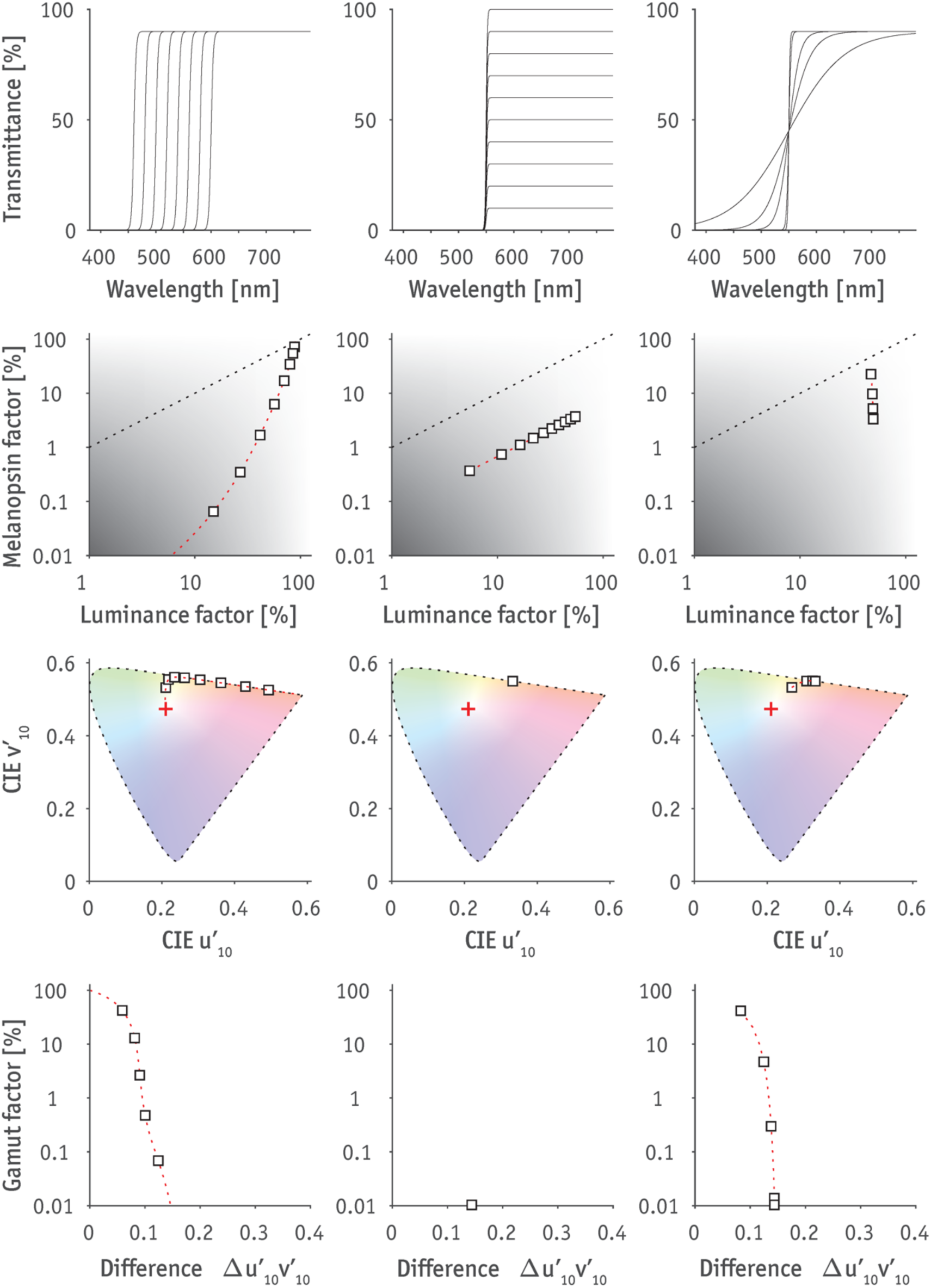
Exploring effects of cut-off filters. Layout follows that of Fig. 3. Spectral transmittance of cut-off filters is analytically modelled using the sigmoid function of the form 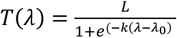, where *L* is the upper asymptote, *k* is the slope, and λ_0_ is the cut-off wavelength. *Left column*: Varying cut-off wavelengths (λ_0_), with *L* = 0.9, *k* = 0.5. *Middle column*: Varying top asymptote L, with λ_0_ *=* 550 nm, *k* = 0.5. *Right column*: Varying slope k, with λ_0_ *=* 560 nm, *L* = 0.9.

Varying the cut-off wavelength systematically affects all four parameters (Fig. 2, *left column*). Moving the cut-off wavelength to longer wavelengths, leads to both more luminance attenuation and more melanopsin attenuation. However, because melanopsin spectral sensitivity has its peak at shorter wavelengths than the luminosity function, the increase in melanopsin attenuation happens at a faster rate than the increase in luminance attenuation. In the colour domain, cut-off filters move the chromaticity towards the spectral locus, with larger deviations with longer cut-off wavelengths.

At a fixed wavelength and slope, varying the asymptote, i.e. the “plateau” and maximum transition systematically effects only luminance and melanopsin, though to the same amount (Fig. 2, *middle column*), producing a line that is parallel to the neutral density locus as one would expect. The chromaticity of filters with varying plateaus is the same. The slope of the transmittance function (Fig. 2, *right column*) at a fixed cut-off wavelength affects largely the attenuation of melanopsin and the change in colour properties, but not so much luminance, though we expect that this will depend on the cut-off wavelength itself.

In this work, we examined 120 “blue-blocking” or “blue-attenuating” filters (either for spectacles or for contact lenses) with respect to their luminance factor, melanopsin factor, colour shift, and gamut factor. These filters were obtained using digital data extraction techniques from a variety of sources, including published graphs from scientific articles, informative patient brochures and manufacturer brochures. We classify these filters in three loose yet intuitive ad-hoc categories: medical filter (n=77), safety filters (n=11) and recreational filters (n=32), which is subdivided into filters for sports (n=12), driving (n=4), visual display unit (VDU) (n=7) use and a catch-all “Other” category (n=8). In this analysis, we are agnostic to the materials that these filters are applied on or produced from, though this is an important consideration for practical use.

In our analysis we find that there is large variability in the visual and non-visual properties of so-called “blue-blocking” or “blue-attenuating” filters.

## Methods

### Data sourcing and extraction

Published graphs of the spectral transmittance for “blue-blocking” and “blue-attenuating” filters were sourced using informal searches on PubMed, Google Scholar, and Google, yielding a total of 120 filters under consideration. Where data were obtained from manufacturers’ or other websites, those websites were submitted to the Internet Archive Wayback Machine for permanent archiving (https://web.archive.org/).

Transmittance spectra were extracted by one operator (author of this study, R.L.) from published graphs using WebPlotDigitzer 4.1^59^, a free visual tool for extraction of data points from published graphs where no tabulated data are available. WebPlotDigitizer has been found to provide rather high levels of reliability^60^. After extraction, data were then interpolated to 1 nm resolution using piecewise cubic hermite interpolating polynomial (PCHIP) interpolation. At wavelengths at which no data was available, the transmittance was set to 0, which has the effect that light at those wavelengths is not factored into the calculation.

### Calculations

For calculation of melanopsin activation, we used the spectral sensitivity recently standardised by the CIE^61^, which assumes a Govardovskii nomogram (λ_max_ = 480 nm) as well as a custom lens function synthesizing different lens transmittance functions^4^. While the standard also recommends functions for the cones^62^, we opted for the CIE 1964 10° observer in this work to maximise compatibility of the colorimetric calculations. This observer prescribes XYZ functions which are converted to the well-known u’_10_v’_10_ diagram. Luminance factor was calculated as the ratio between the luminance of the light seen with or without the filter incorporated as a wavelength-wise multiplication. The same procedure was used for the melanopsin factor calculation. Colour differences were calculated using the Euclidiean distance between two points in the diagram u’_10_v’_10_. The colour gamut was calculated using a set of 99 representative reflectances^63^. The gamut was calculated using the ratio of the area of the convex hulls of these 99 reflectance functions seen with and without the filter. The different effects are visualised in Figure 1. For all calculations, we assumed a spectrally-neutral equal energy white (EEW), which has equal power at all wavelengths. We realise that this illuminant is at best a hypothetical one, but suggest that it is a suitable one for the purpose of objectively comparing the filtering properties of different filters without recourse to a specific class of illuminants.

### Data and software availability

All tabulated transmittance spectra and code to produce the unedited versions in the figures in this report are available on GitHub (https://github.com/spitschan/bbFilter).

## Results

### Large diversity of filters

Across all categories, the spectral transmittances of the “blue-blocking” and “blue-attenuating” filters are rather diverse (Fig. 3, *Row 1*). This diversity is exhibited in whether the filter is a cut-off, notch or has some other filter shape. The cut-off filters differ largely in the wavelength at which they transmit 50% of light. Another differing factor is the “plateau” of transmission at wavelengths longer than the cut-off wavelength.

**Figure 3:**
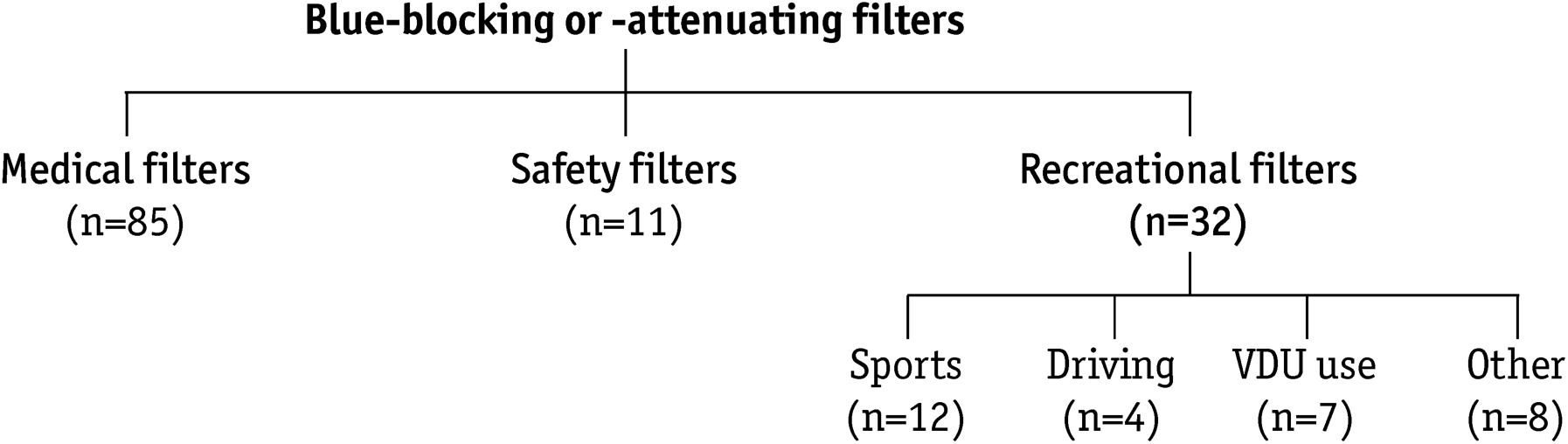
Ad-hoc taxonomy for the examined filters.

**Figure 4:**
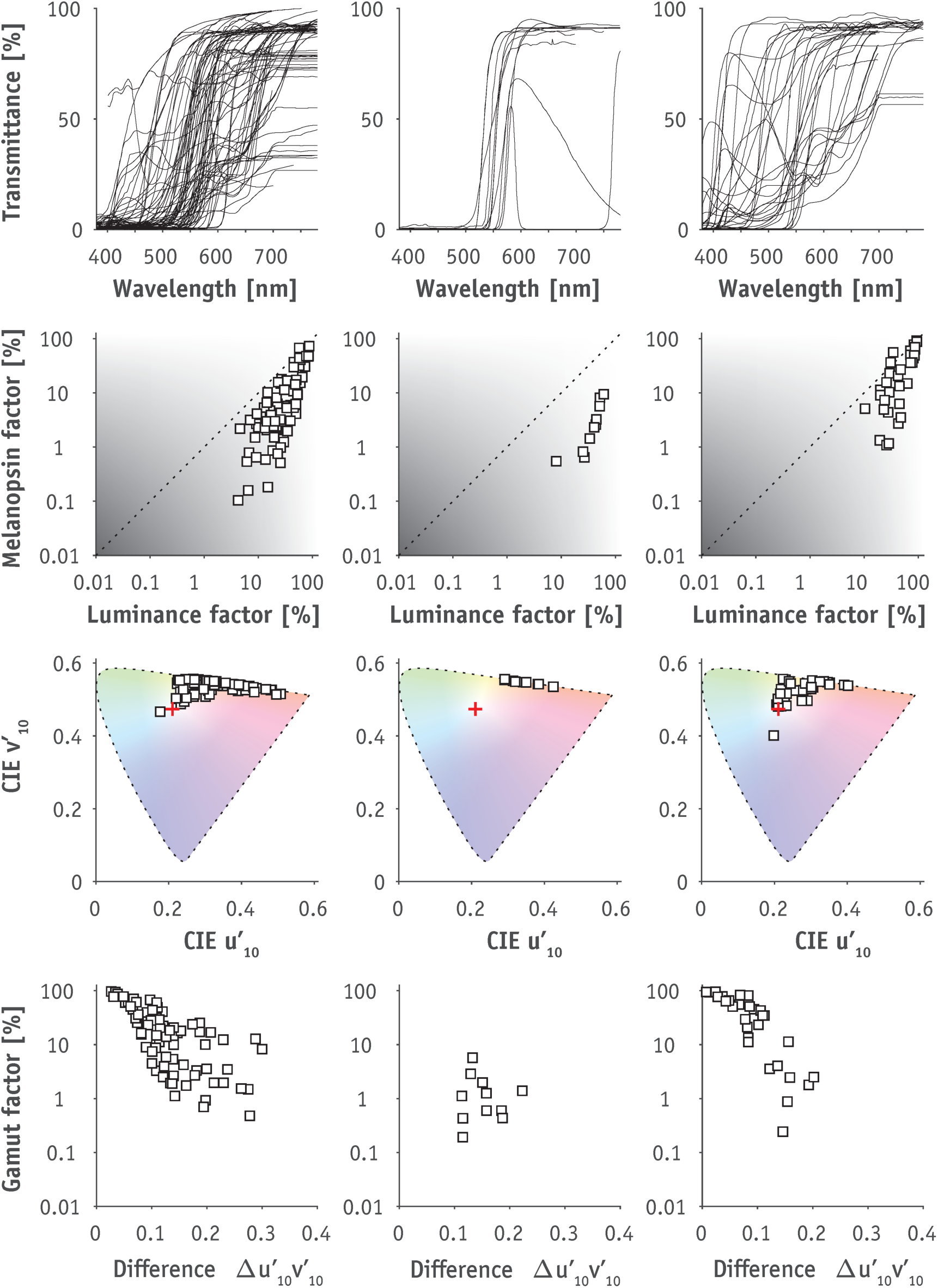
Visual and non-visual properties of “blue-blocking” and “blue-attenuating” optical filters. *Columns correspond to the three main filter categories we identified (Fig. 2). Row 1: Spectral transmittances of filters. Row 2: Luminance factor vs. melanopsin factor. Dashed line indicates equal reduction of luminance and melanopsin, as would be the case with a spectrally uniform neutral density (ND) filter. Row 3: Chromaticity diagram. Red plus indicates chromaticity of equal-energy white (EEW) and white squares indicate chromaticities of EEW seen through the respective filters. Row 4: Colour shift vs. gamut factor. See Introduction and Fig. 1 for explanation.*

Nearly all filters show more relative melanopsin attenuation than luminance attenuation (*Fig. 3, Row 2*). This is evidenced by the fact that all data points lie under the “diagonal”, which corresponds to a spectrally flat filter which decreases the activation of all photopigments by the same amount. It appears as though there is a “fan” effect in the relationship between luminance and melanopsin attenuation, bounded by the neutral density diagonal at the upper end.

Nearly all filters show a move of the chromaticity of the EEW white point towards the spectral locus, which is given by the edge of the chromaticity diagram. This locus corresponds to the chromaticity of monochromatic and therefore highly saturated lights. Interestingly, there is again a “fan” effect in the relationship between luminance and chromaticity. The intuition here is that the colour matching functions used to calculate chromaticity of the white point include the luminosity function, and therefore, these effects are not independent except under special conditions.

Regarding the colour rendering properties of the filters quantified using the gamut attenuation, we find no clear pattern. We find that some filters severely reduce the colour gamut of the worlds seen with the filter, reducing it to only <1% of the gamut seen with the filter. This is obviously correlated with the shift of the filter towards the spectral locus, as all surfaces seen under monochromatic illumination appear to only change in lightness, and not colour (because the surfaces can only reflect the light that is there). As would be expected, larger colour differences (i.e. shift of the colour towards the spectral locus) translate to a reduced colour gamut.

## Discussion

### General discussion

In our analysis we find that there is large variability in the visual and non-visual properties of so-called “blue-blocking” or “blue-attenuating” filters. While the spectral transmittance of a filter necessarily needs to be known for quantifying the four outcomes we investigated here, it in itself is not of any use as to determine a filter’s effect on the retina. Rather than simply (or not, in some cases) providing transmittance spectra in graphical or tabulated form, future studies should quantify the effect of a filter using the metrics described here.

For many of the applications of filters manipulating short-wavelength light reaching the retina, we still lack a mechanistic understanding of how the different photoreceptors contribute to the effects. For example, at present (2019), we do not know how cones and rods contribute to the basic physiological regulation of melatonin secretion by light. A retinally referenced, or “physiologically relevant” framework to quantify effects of a filters is the first step in using optical filters for developing such mechanistic understanding.

### Pupil size effects

The reduction of illumination at cornea by optical filters reduces the retinal illuminance, i.e. the incident light on the retinal surface. However, retinal illuminance itself is regulated by the pupil area. With a reduction in light intensity, the pupil area becomes bigger. The dynamic range, however, is rather limited, with a maximum possible modulation of retinal illuminance just by pupil size by a factor of ∼16 (between a maximally constricted 2 mm pupil and a maximally dilated 8 mm pupil). We investigated the effect of reducing the corneal illuminance with optical filters on the effective retinal illuminance using the unified Watson & Yellott model^64^ (Fig. 5). We assumed a field size of 150°, a 32-year old observer, as well as binocular stimulation as an approximation to real world viewing conditions, and investigated how the predicted retinal illuminance depends on the luminance of the viewed stimulus with and without neutral density filters of varying densities (ND1.0, ND2.0 and ND3.0). With the near-parallel lines of log luminance vs. log retinal illuminance, it can be seen that the primary determinant of retinal illuminance is luminance “seen” through the filter.

**Figure 5:**
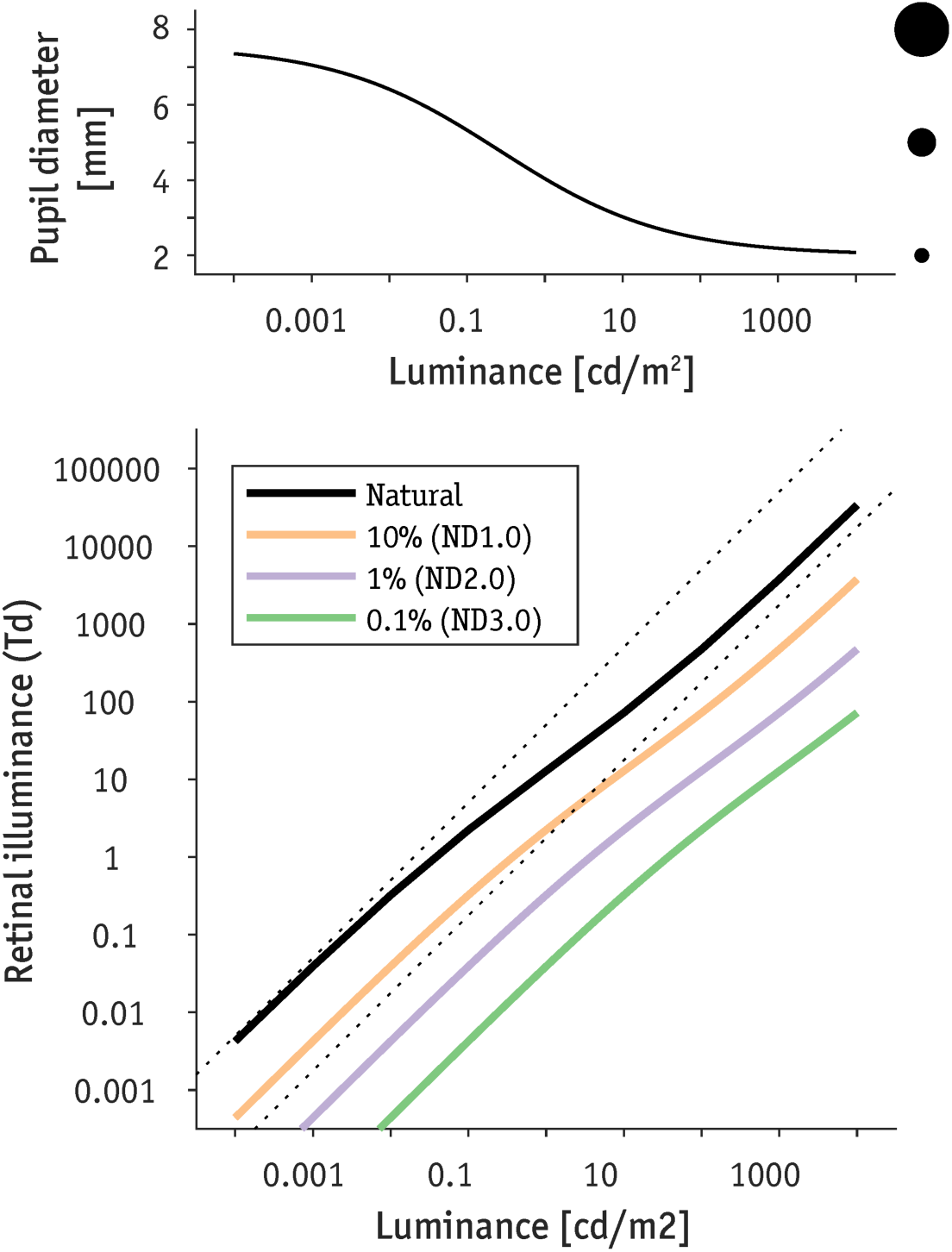
Pupil size effects of neutral-density filters. *Top panel: Predicted pupil size for a 32-year old observer viewing a 150° field at varying luminances with both eyes. Bottom panel: Predicted retinal illuminance (luminance × pupil area) when viewed either through the natural pupil (dashed lines indicate maximum retinal illuminance given maximum difference in pupil size), or through spectrally uniform filters of varying transmittance (ND1.0, ND2.0, ND3.0).*

Chung and Pease ^14^ note that at equivalent luminance, “yellow” filters lead to larger pupils than neutral-density filters. This is consistent with the view that melanopsin activation, which significantly drives steady-state pupil size in humans^65,66^, is severely reduced under short-wavelength filters. Under most real-world conditions, the correlation between luminance and melanopsin activation is probably well-constrained, except with chromatic filters and experimental conditions in which the decoupling can be lead to up to a three-fold difference in melanopsin activation with no or little nominal difference in luminance^67,68^. Importantly, optical quality of the retinal image depends on pupil size^69^, which needs to be factored into filter assessments.

### Digital data extraction

This study relied on digital data extraction from published graphs. This was necessary because spectral transmittance data are rarely available in digital or tabulated form, even though data storage as supplementary material is often available at scientific journals at the time of writing this article (2019). As shown in this article, this need not be a limitation since data can be extracted from graphs and be subjected to rigorous novel or reanalyses. A limitation of our data-driven approach is that tabulated spectra given by manufacturers are here treated as *bona fide* measurements of transmittance spectra. The process of data extraction may also lead to small inaccuracies in the tabulated transmittance spectra. Optical effects such as scattering or polarisation and other higher-order effects are not captured by the simple filter model that we have applied here.

### Other considerations

In addition to filters affecting the retinal stimulus, light of different colours may also have psychological or higher-order effects^70^. Another effect worth considering is stigma towards wearers of coloured spectacles^71^. Here, we only considered the visual and non-visual properties of different optical filters, though these effects will need to be factored into the feasibility of using specific optical filters for a desired physiological or psychological effect.

## Conclusion

We find large diversity in the visual and non-visual properties of different “blue-blocking” or “blue-attenuating” filters. We propose that to evaluate the effect of a given optical filter, the spectral transmittance is only the first step in characterising the effect of a filter on the illumination at the eye and suggest a retinally referenced framework to quantify these effects, incorporating the attenuation of luminance, the attenuation of melanopsin activation, shifts in colour, and reduction of colour gamut.

## Tables

**Table 1.**
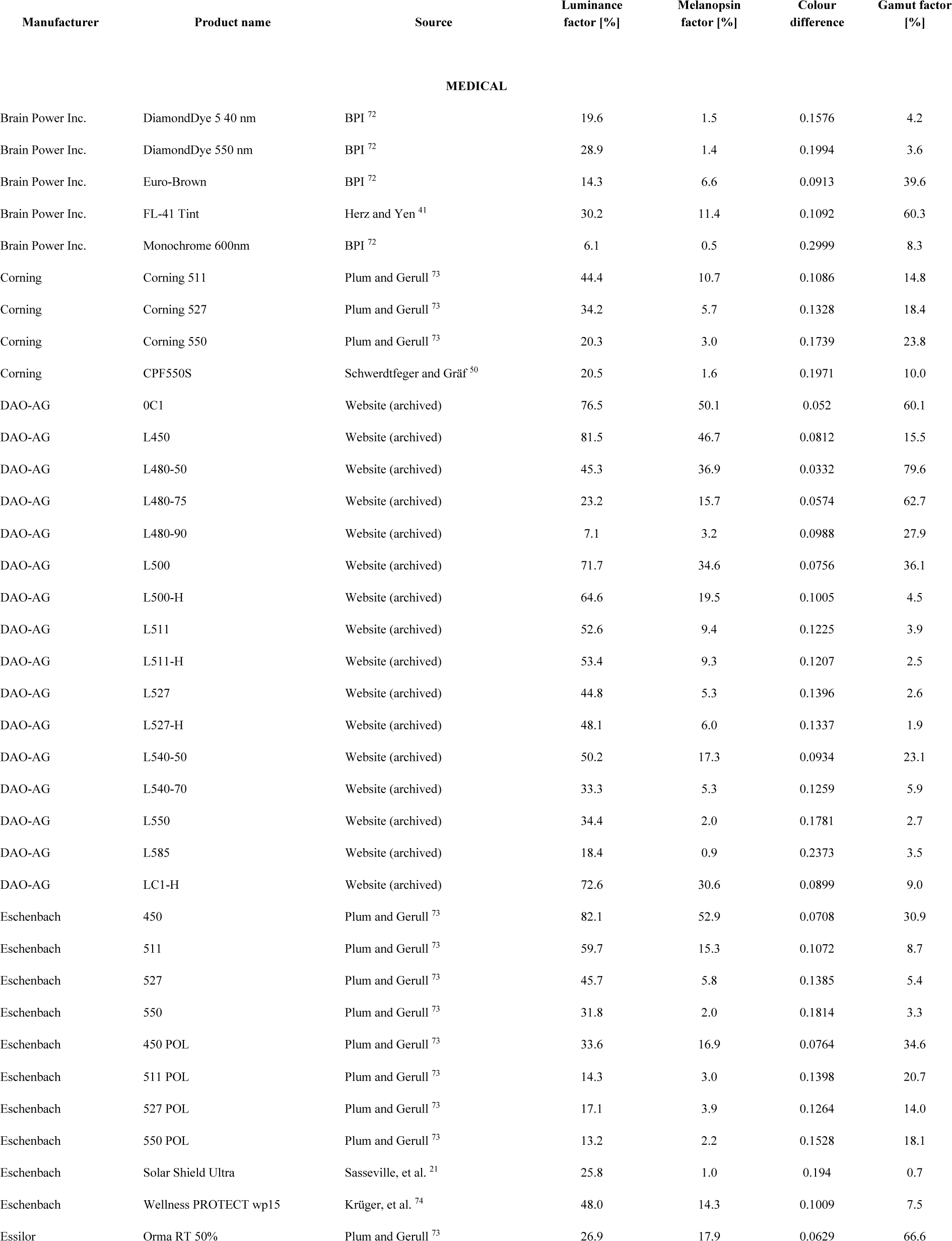
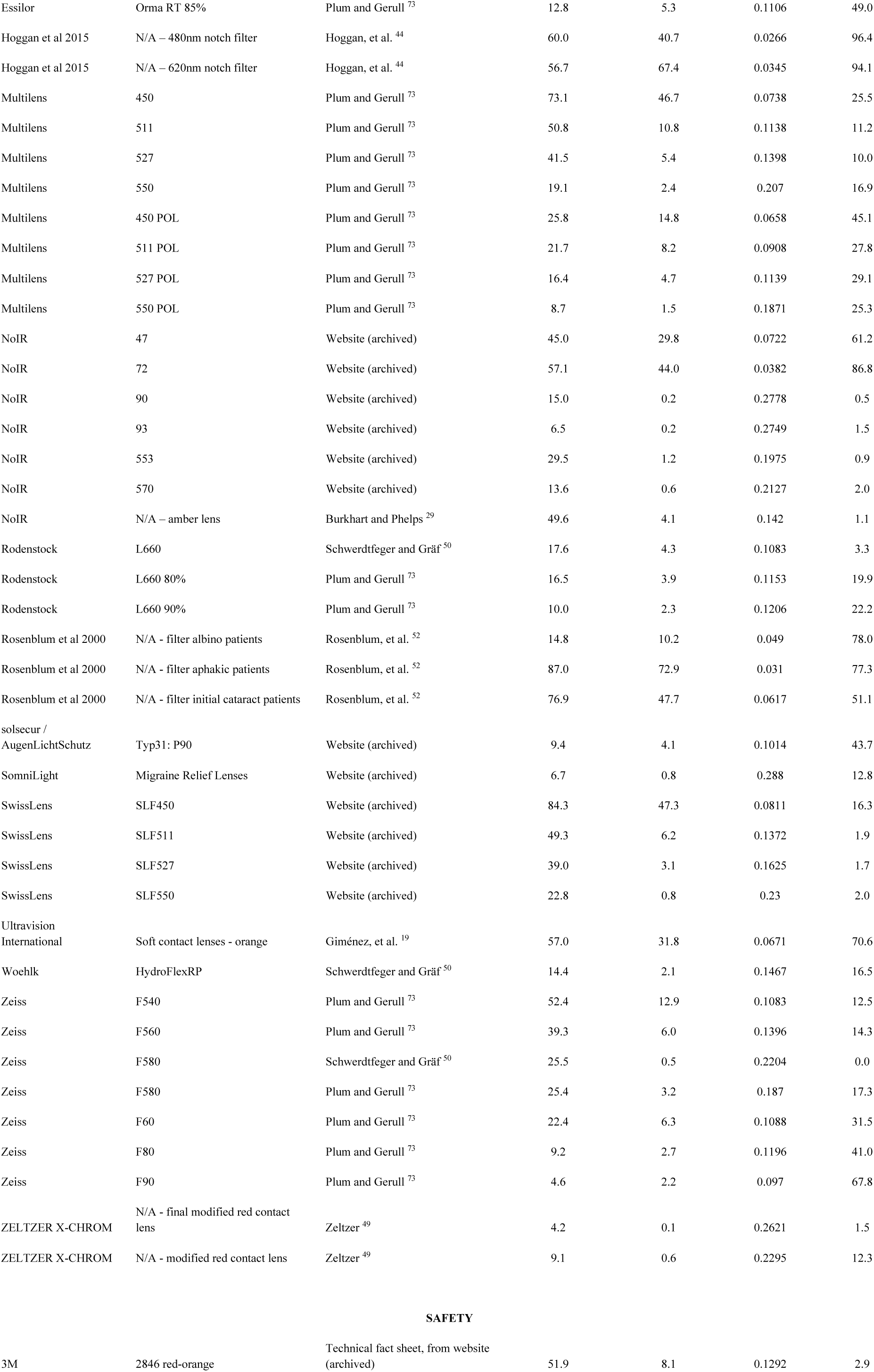
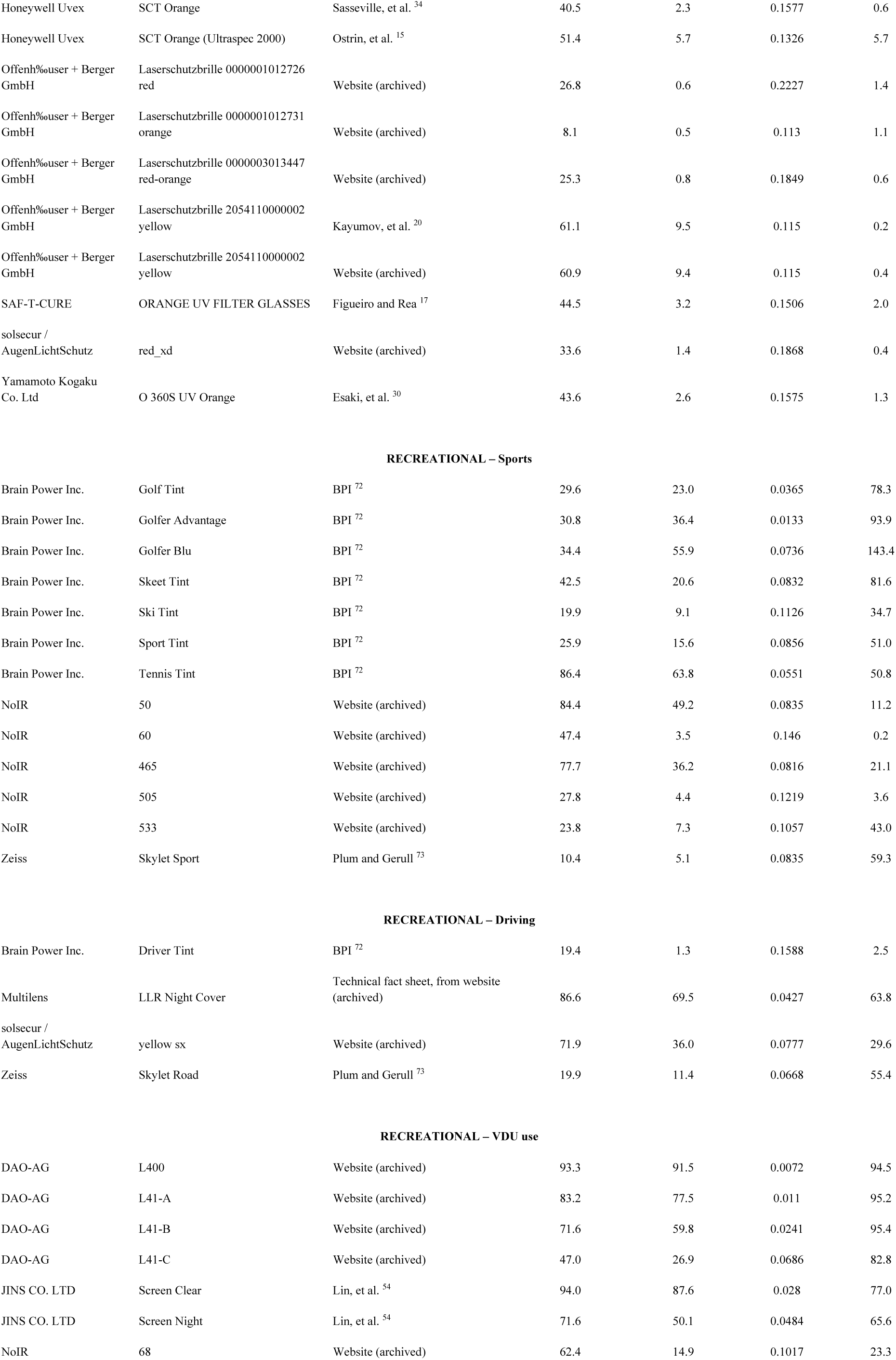
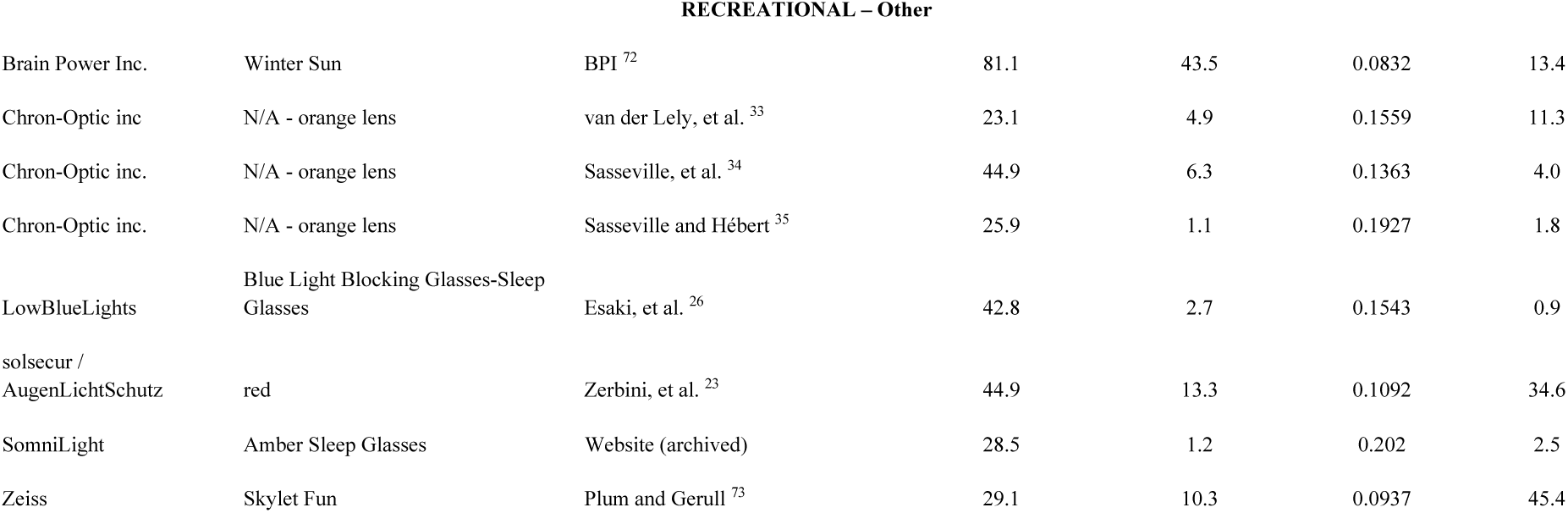

## Acknowledgements

M.S. is supported by a Sir Henry Wellcome Trust Fellowship (Wellcome Trust 204686/Z/16/Z) and a Junior Resaerch Fellowship from Linare College, University of Oxford.

